# Senolytic treatment depletes microglia and decreases severity of experimental autoimmune encephalomyelitis

**DOI:** 10.1101/2024.02.05.579017

**Authors:** Sienna Sage Drake, Aliyah Zaman, Christine Gianfelice, Elizabeth M.-L. Hua, Kali Heale, Elia Afanasiev, Jo Anne Stratton, Alyson Fournier

**Affiliations:** Montreal Neurological Institute, McGill University, Montreal, Quebec, Canada

## Abstract

The role of senescence in disease contexts is complex, however there is considerable evidence that depletion of senescent cells improves outcomes in a variety of contexts particularly related to aging, cognition, and neurodegeneration. Here, the effect of a bioinformatically-rationalized senolytic was tested in the experimental autoimmune encephalomyelitis (EAE) mouse model of multiple sclerosis (MS). Single-cell analysis from brain tissue isolated from mice subjected to EAE identified microglia with a strong senescence signature including the presence of BCL2-family member transcripts. Cells expressing Bcl2l1 had higher expression of pro-inflammatory and senescence genes than their negative counterparts in EAE, suggesting they may exacerbate inflammation. Notably, in human single-nucleus sequencing from MS, BCL2L1 positive microglia were strongly enriched in lesions with active inflammatory pathology, and likewise demonstrated increased expression of immune related genes suggesting they may contribute to the active lesion pathology and tissue damage in chronic active lesions. Employing a small molecule BCL2 inhibitor, Navitoclax (ABT-263), significantly reduced the presence of microglia in the EAE spinal cord, suggesting that these cells can be targeted by senolytic treatment. ABT-263 treatment had a profound effect on EAE mice, decreasing motor symptom severity, improving visual acuity, promoting neuronal survival, and decreasing white matter inflammation. Together, these results provide evidence to support the idea that senescent glia may exacerbate inflammation resulting in negative outcomes in neuroinflammatory disease and that removing them may ameliorate disease.

## Introduction

Multiple sclerosis (MS) is a complex immune and neurodegenerative disease. The classic pathological feature of MS is the inflammatory demyelinating lesion which consists of myelin sheath degeneration and axonal injury. Extravasation and activation of T-cells, B-cells, and macrophages in the central nervous system (CNS), alongside gliosis of resident astrocytes and microglia contribute to demyelination, axonal injury, and neuronal and oligodendrocyte cell death^1^. Transcriptional studies of CNS glial cells have identified numerous dysregulated pathways, including complement signaling, interferon signaling, and antigen presentation in astrocytes, microglia, and oligodendrocytes^2^.

Of the CNS glial cells, microglia are particularly implicated in disease development and progression. Microglia are a main contributor to the pathology of chronic active or smouldering lesions, which are characterized by the presence of a rim of reactive microglia and macrophages that contribute to gradually expanding tissue injury^3,4^. Chronic active lesions and microglia activation and reactivity are also associated with MS disease progression and severity, even in the absence of new or ongoing peripheral immune cell invasion and inflammation^4,5^. Reactive microglia can produce high amounts of the cytokine TNFα which is toxic to oligodendrocytes thus likely contributing to demyelination and oligodendrocyte cell death^4^. Treatments that deplete microglia are likewise effective at reducing disease severity in rodent models of MS^4^. However, microglial cell states in the CNS are highly complex, being influenced by their microenvironment, the molecules they express, and their function^6^. In some contexts, microglia promote tissue injury and inflammation, while in other contexts, microglia assume a phagocytic role in clearing myelin debris to promote remyelination and repair^4,7–9^. The identification of strategies to specifically target injurious microglia could have an important role in reducing disease activity within the CNS and in tackling progression in MS.

The continuum of microglial states from homeostatic to injurious exists upon the axes of healthy to injured and young to old^10^. As a dividing cell population, microglia are susceptible to cellular senescence, and senescent microglia can perpetuate CNS inflammation and injury^11^. Importantly, aged microglia demonstrate decreased turnover and limited myelin debris phagocytosis suggesting impaired homeostatic functions^12,13^. Like an ouroboros, CNS injury and inflammation can likewise accelerate microglial senescence. This has been particularly demonstrated in models of traumatic brain injury which induce a senescent profile in microglia; depleting these microglia subsequently improves outcomes related to cognition and tissue repair^14–16^. Further evidence for the relationship between aging and inflammation in microglia comes from an elegant study which conducted adoptive transfer EAE (AT-EAE) in young and middle-aged mice^17^. Here, AT-EAE in middle aged mice produces a significantly more severe and progressive disease course. Additionally, single-cell RNA-sequencing demonstrated that, compared to microglia from young AT-EAE mice, microglia from middle aged AT-EAE mice downregulate homeostatic marker genes, and upregulate genes related to proinflammatory interferon signaling, antigen presentation, and proteasome assembly^17^.

Since biological age is correlated with conversion from relapsing-remitting to progressive MS, it is possible that disease progression without peripheral immune involvement is driven in part by the persistent accumulation of proinflammatory, senescent cells^18^. Senescent cells canonically upregulate anti-apoptotic genes including the BCL2 family genes, enabling them to evade cell death and persist longer in tissue, exacerbating inflammation and tissue injury^19^. Thus, drugs that can specifically target senescent cells may interrupt this vicious cycle. Senolytics are one such class of drugs that induce cell death in senescent cells and penetrate the CNS to eliminate senescent glia^16,20,21^. One notable study found senolytic treatment prevented the upregulation of senescence genes and attenuated *tau* phosphorylation in a mouse model of Alzheimer’s disease^22^.

Given the strong relationship between MS progression, aging, and microglia, this study aimed to investigate whether microglia in EAE and in MS exhibit markers of senescence, and whether they might be eliminated by senolytic treatment. Here, analysis of single-cell RNA-sequencing data from EAE and control mice establishes a proinflammatory and senescent microglial transcriptome in the disease. Assessment specifically of cells expressing the anti-apoptotic gene *Bcl2l1* demonstrates that these cells exhibit even higher levels of proinflammatory and senescence-associated genes in EAE compared to their *Bcl2l1* negative counterparts. In human MS single-nucleus sequencing, *BCL2L1* was enriched in microglia in lesions with chronic active pathology, and *BCL2L1* positive cells demonstrated concurrent upregulation of genes relevant to senescence and inflammation. Navitoclax (ABT-263), a small molecule BCL-2 family inhibitor and senolytic drug, led to significant depletion of microglia in EAE mice, reduced motor symptom severity, improved neuronal survival, and diminished markers of inflammation in the white matter. Together, this study provides evidence that senolytic therapy through BCL2 inhibition may be a promising avenue to reduce CNS inflammation and prevent disease worsening in MS by elimination of senescent microglia.

## Results

### Analysis of single-cell RNA sequencing and immunohistochemistry of EAE microglia implicates cellular senescence

To characterize microglial gene expression in the context of pathological inflammation, single cell RNA-sequencing data from EAE and control mice was analyzed, and microglia were identified based on annotations from the published dataset^23^ (Fig. 1A). In this cell cluster, high expression in of microglial transcripts such as Aif1, P2ry12, Csf1r, and Cx3cr1 confirmed the microglial identity (Fig. 1B-E). Microglial gene expression was aggregated across biological replicates (n=3 Control, n=3 EAE) and differential gene expression analysis followed by KEGG pathway level analysis was conducted. KEGG pathways strongly prioritized pathways associated with neurodegeneration, including “Alzheimer’s disease” and “Parkinson’s disease”. Other biological pathways pointed towards cellular injury and senescence, including “Oxidative phosphorylation”, “Cell cycle”, “Cellular senescence”, and “Necroptosis”. Further pathways highlighted were related to DNA damage and repair, and proinflammatory signaling cascades including “Antigen processing and presentation”, “TNF signaling pathway”, and “NF-kappa B signaling pathway” (Fig. 1F). Thus, pathway level analysis suggests EAE microglia have a proinflammatory, neurodegenerative, and senescent phenotype. A canonical feature of senescent cells is the upregulation of anti-apoptotic Bcl2 family genes, which can be targeted by a class of senolytic drugs that function through their inhibition^19^. These include *Bcl2/*BCL2, *Bcl2l1*/BCL2-xL, and *Bcl2l2*/BCL2-w. Given the strong inflammatory and senescent transcriptional phenotype in EAE microglia, the three Bcl2 family genes were assessed to evaluate whether senescent EAE microglia might be targetable by BCL2 inhibiting senolytics. Indeed, all markers were higher in EAE compared to control cells, with *Bcl2l1* notably being significantly upregulated and detected in over 15% of EAE microglia (Fig. 1G). Given its significant upregulation in EAE and higher proportion of positive cells, *Bcl2l1* was subsequently used to partition EAE microglia, and differential gene expression between *Bcl2l1* positive and *Bcl2l1* negative EAE microglia was conducted. Here, EAE microglia expressing *Bcl2l1* showed significantly higher expression of many genes related to proinflammatory processes including *Tnf*, *Tlr2*, and *Icam1* (Fig. 1H). Additional upregulated genes associated with aging, senescence, and inflammation notably upregulated in *Bcl2l1*+ EAE microglia included *Ccl2* and *Cdkn1a*. To better assess the specific contribution of senescence related genes to EAE microglia, genes from a curated senescence gene set were visualized based on significance and differential expression^24^. In EAE microglia, 41 of the 101 detected genes from this gene-set were dysregulated, with 33 being significantly upregulated in EAE (Fig. 1I). These 41 genes were then assessed in the *Bcl2l1* positive EAE microglia. Here, compared to *Bcl2l1* negative EAE microglia, *Bcl2l1* positive EAE microglia showed further significant upregulation of 20 of the 41 genes, supporting the hypothesis that *Bcl2l1* positive microglia exhibit higher levels of senescence associated genes than their negative counterparts (Fig. 1J). To further characterize the *Bcl2l1* positive microglial subset in EAE, GO term analysis of significantly upregulated genes was conducted. This analysis revealed a strong inflammatory and cellular stress signature, with terms such as “activation of innate immune response”, “regulation of autophagy”, “response to ER stress”, “canonical NFkb signal transduction”, and “lymphocyte activation involved in immune response” (Fig. 1I). Together, these data suggest that EAE microglia display a strong inflammatory and senescent signature. The subset of microglia that express anti-apoptotic Bcl2 family genes express higher levels of proinflammatory and senescence genes than their negative counterparts, and display broad dysregulation of pathways related to inflammation, mRNA processing, autophagy, and chromatin remodeling. To validate that there are in fact senescent microglia in the white matter, IHC of EAE optic nerve was conducted for BCL2, IBA1, and senescence associated beta-galactosidase (SABG) (Fig. 2A-D). Confocal imaging demonstrated the unequivocal presence of numerous cells triply labeled for BCL2, IBA1, and SABG. Indeed, BCL2 staining particularly labeled cells with a microglial like morphology (Fig. 2E-I). Together, these data provide evidence that microglia demonstrate a senescent profile combined with expression of BCL2.

**Figure 1.**
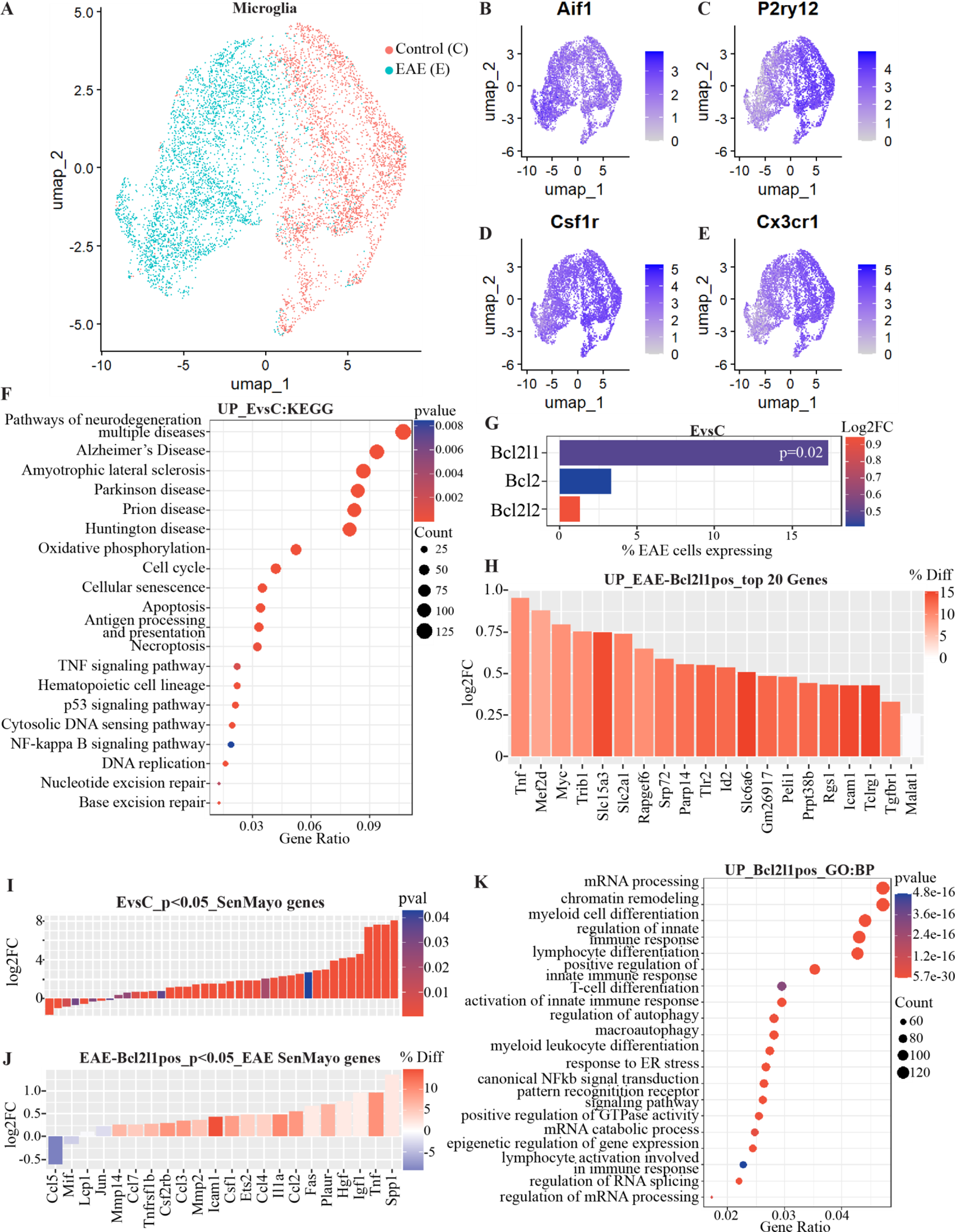
Single-cell RNA-seq of EAE microglia implicates senescence and identifies inflammatory subpopulation expressing ABT-263 targets. A-E) Microglial cells from Fournier et al. 2023 from EAE and Control mice, as confirmed by expression in the cluster of microglial specific genes Aif1 (B), P2ry12 (C), Csf1r (D), and Cx3cr1 (E). F) Kegg pathway enrichment for genes upregulated in EAE microglial demonstrates an enrichment for immune signaling pathways, cellular senescence, and neurodegenerative diseases. G) Percent of EAE microglia expressing various targets of ABT-263 and log2FC between EAE and control microglia. H) Top 20 upregulated genes in Bcl2l1 positive EAE microglia compared to Bcl2l1 negative EAE microglia, colour represents the difference in % positive cells between Bcl2l1+ population and Bcl2l1-population. I) Genes significantly dysregulated in EAE vs Control microglia belonging to the SenMayo dataset (n=41 genes, up=33,down=8, gene labels removed for clarity of figure). J) Dysregulation of the 41 SenMayo genes dysregulated in EAE in Bcl2l1 positive EAE microglia, colour represents the difference in % positive cells between Bcl2l1+ and Bcl2l1-populations, positive number/red indicates higher percent positive cells in Bcl2l1+ population, negative number/blue indicates higher percent positive cells in Bcl2l1-population (n=22 genes, Log2FC up=20, Log2FC down=2). K) GO biological process pathway enrichment for upregulated genes in Bcl2l1 positive EAE microglia demonstrating top enriched processes.

**Figure 2.**
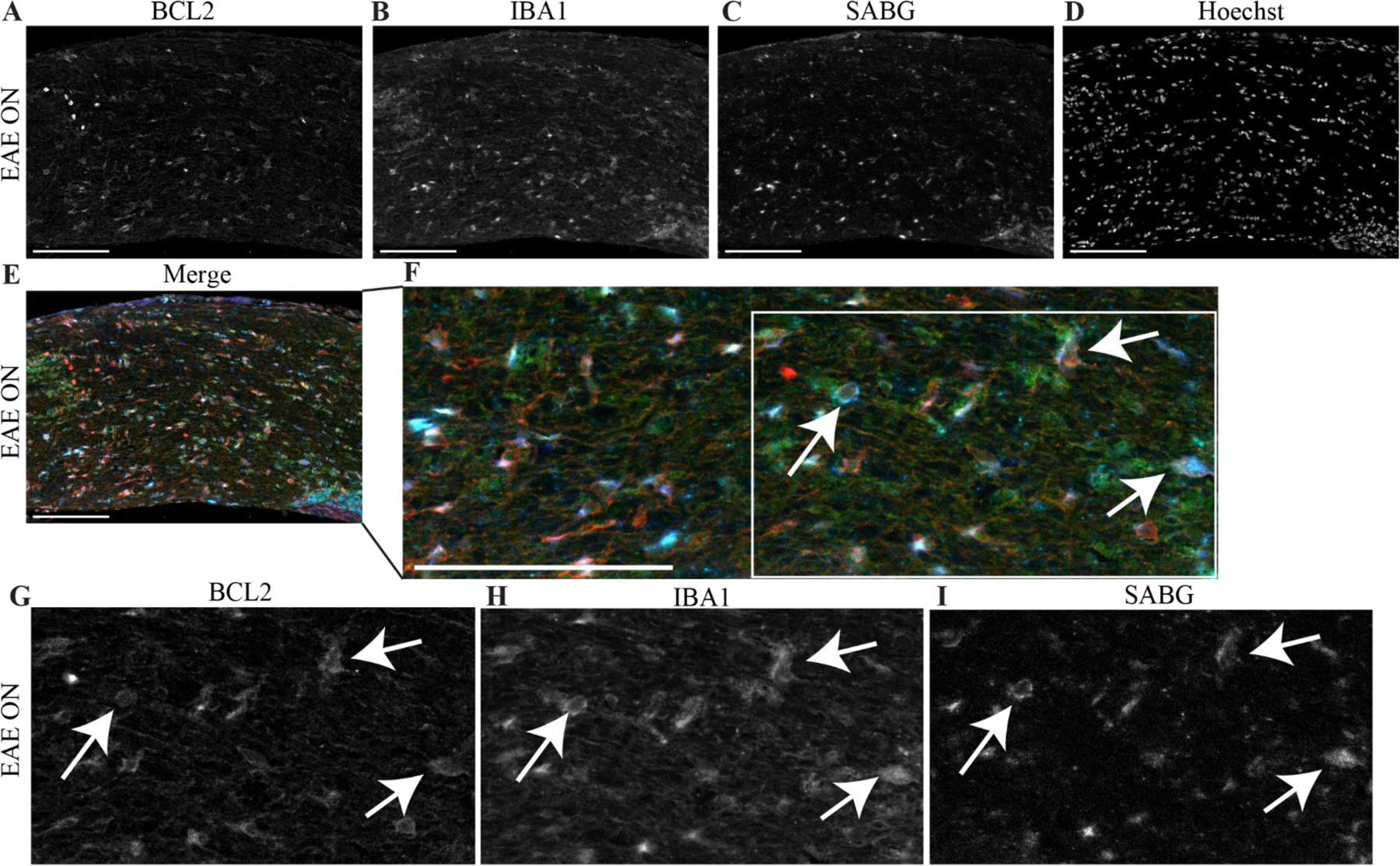
EAE white matter microglia express BCL2 and senescence associated beta-galactosidase. **A-D)** Example of EAE optic nerve immunostained for BCL2 (A), IBA1 (B), senescence-associated beta galactosidase (C), and Hoechst (D). **E,F)** merged image of BCL2 (red), IBA1 (green), and senescence associated beta galactosidase (blue) with zoomed image (F) demonstrating triple positive labeled cells (arrows) throughout the optic nerve. Scale bar = 100 um. **G-I)** Zoomed images of individual channels for BCL2 (G), IBA1 (H), and SABG (I) to demonstrate positive cellular staining.

### Human MS microglia display enrichment of *BCL2* and *BCL2L1* in chronic active lesions

As chronic active lesions are strongly associated with disease severity, this study first aimed to evaluate the microglial transcriptome in chronic active lesions. Two single-nucleus sequencing datasets of human MS patient tissue were acquired from the GEO^25,26^. The first dataset consisted of cells from healthy control tissue, MS patients with chronic active lesion pathology, and MS patients with chronic inactive lesion pathology^25^ (Fig. 3A). The published cell type annotations were used to identify the microglial cell cluster, which demonstrated enrichment for microglial genes (Fig. 3B). Notably, comparing the microglia cluster across the annotated pathologies demonstrated an enrichment for *BCL2L1* expression in chronic active samples, but not from control or chronic inactive samples (Fig. 3C). The second dataset was processed similarly, however here annotations from the original publication labeled a group of cell clusters as ‘immune’. Cluster 7 in this immune annotation was used in downstream analyses due to its stronger enrichment for microglial genes compared to other immune labeled clusters (Fig.3D,E). Again, *BCL2L1* expression was the highest and in the most cells in microglial cells coming from chronic active labeled lesion pathology, suggesting that these cells may be contributing to ongoing inflammatory activity and tissue injury in this type of lesion (Fig. 3F). Analysis of differentially expressed genes between the *BCL2L1*+ and *BCL2L1*-cells within this dataset identified the upregulation of several genes related to apoptosis including *BCL2* and *BIN1* (Fig. 3G-I). Genes related to cellular growth signaling including *PI4KA* and *RPTOR*, were likewise upregulated (Fig. 3J,L). Additional upregulated genes were related to inflammation, including *PTPN1* which is known to regulate the neuroinflammatory response in microglia, *NFKB1* which is an integral component of the NFKB signaling pathway, *TNFSRF1A* which is a TNF receptor involved in transducing TNF signaling cascades and a GWAS gene for MS susceptibility, *ILRAP* which is involved in IL1 signaling and microglial activation, and *CADM1*, a cell adhesion molecule that plays a role in NFKB signaling during virus infection^27–33^(Fig. 3K-P). Together, these data suggest that MS microglia are particularly enriched for *BCL2L1* in chronic active lesions, and *BCL2L1* positive human microglia, like EAE microglia, have an altered gene profile with contribution of known proinflammatory genes.

**Figure 3.**
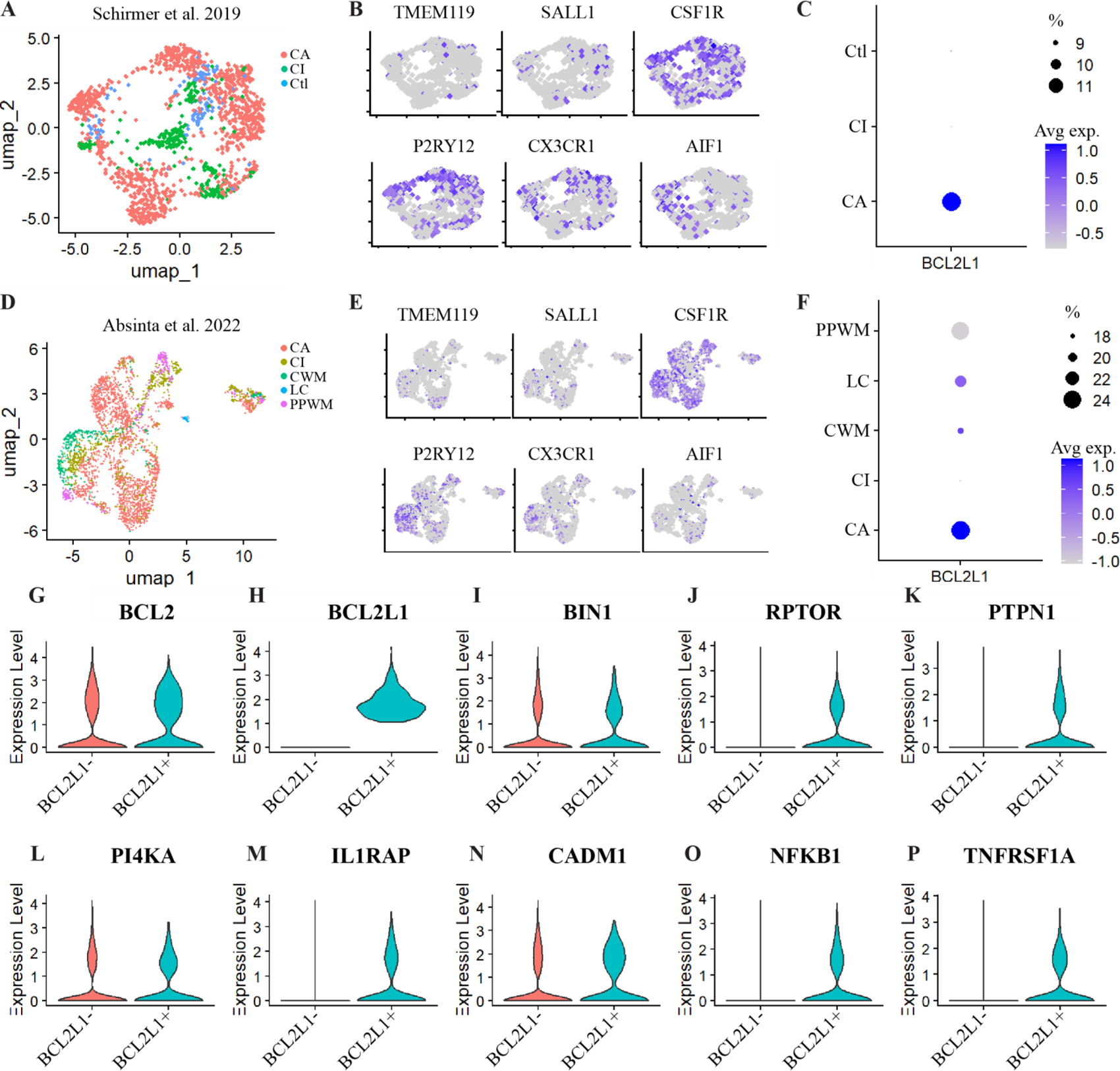
Single-nucleus RNA-seq of human MS microglia demonstrate enrichment for *BCL2L1* genes in chronic active lesions and dysregulation of immune genes in BCL2L1+ cells. **A)** Microglia from Schirmer et al. 2019 clustered based on microglial annotation accompanying the original published manuscript across control samples (Ctl), MS tissue with chronic active lesion pathology (CA) and chronic inactive lesion pathology (CI). **B)** Microglial marker gene expression in the microglia annotated cluster. **C)** Expression and percent cells expressing BCL2L1 across different conditions. **D)** Microglia from Absinta et al. 2022, obtained from ‘immune’ annotation provided by original publication and then subset on cluster 7 based on highest presence of microglial markers across different lesion pathologies including chronic active lesion edge (CA), chronic inactive lesion edge (CI), lesion core (LC), control non-lesion white matter (CWM), and periplaque white matter (PPWM). **E)** Microglial marker gene expression. **F)** Expression and percent cells expressing BCL2L1 across the different lesion classifications. **G-P)** Violin plots of select upregulated genes comparing BCL2L1 positive and BCL2L1 negative microglia in the Absinta et al dataset, demonstrating upregulation of genes related to neuroinflammation.

### Senolytic treatment depletes microglia in EAE spinal cord

Given the presence of senescent microglia in EAE and MS with expression of Bcl2 family genes, the small molecule BCL2 inhibitor, Navitoclax (ABT-263), was administered to EAE mice and immune cell flow cytometry conducted on spinal cord immune cell isolates (Fig. 4A). ABT-263 was selected as it is a high affinity inhibitor for all three members of the BCL2 family proteins (BCL2; *Bcl2*, BCL-xL; *Bcl2l1*, and BCL-w; *Bcl2l2*), is a known senolytic, and penetrates the CNS^34,35^. Drug administration was conducted daily beginning at EAE symptom onset to not interfere with disease induction, and flow cytometry analysis was conducted at peak disease at least 3 days post treatment start. Cells were gated to enable analysis of percent population of viable single cells (Fig. 4B-D). CD45 and Cd11b were used to identify immune cells and differentiate the innate immune cells from lymphocytes (Fig. 4E-F). Microglia were identified based on their medium expression of CD45, and further differentiated based on low expression of Ly6G and Ly6C (Fig. 4F,H). CD45 high Cd11b positive cells were similarly grouped based on Ly6G and Ly6C expression, corresponding broadly to monocytes/macrophages (Ly6C-,Ly6G-), neutrophils (Ly6G+), and proinflammatory monocytes (Ly6C+) (Fig. 4G). CD45 positive CD11b negative lymphocytes were stratified by CD4 and CD8 expression to identify the two major classes of T-cells (Fig. 4I). Within the macrophage/monocyte/neutrophil populations, there were no significant changes in cell number, although the Ly6C-Ly6G-monocyte/macrophage population tended towards a decrease (Fig. 4J-M). Importantly, in the microglial populations, there was a small but significant decrease in the percent of the viable cell population (Fig. 4N,O). Neither population of lymphocytes demonstrated a change in percentage of the cell population (Fig. 4P,Q). Thus, these data demonstrate that senolytic treatment with a small molecule BCL2 inhibitor specifically depletes microglia in EAE spinal cord seemingly without altering other major cell types of the immune system.

**Figure 4.**
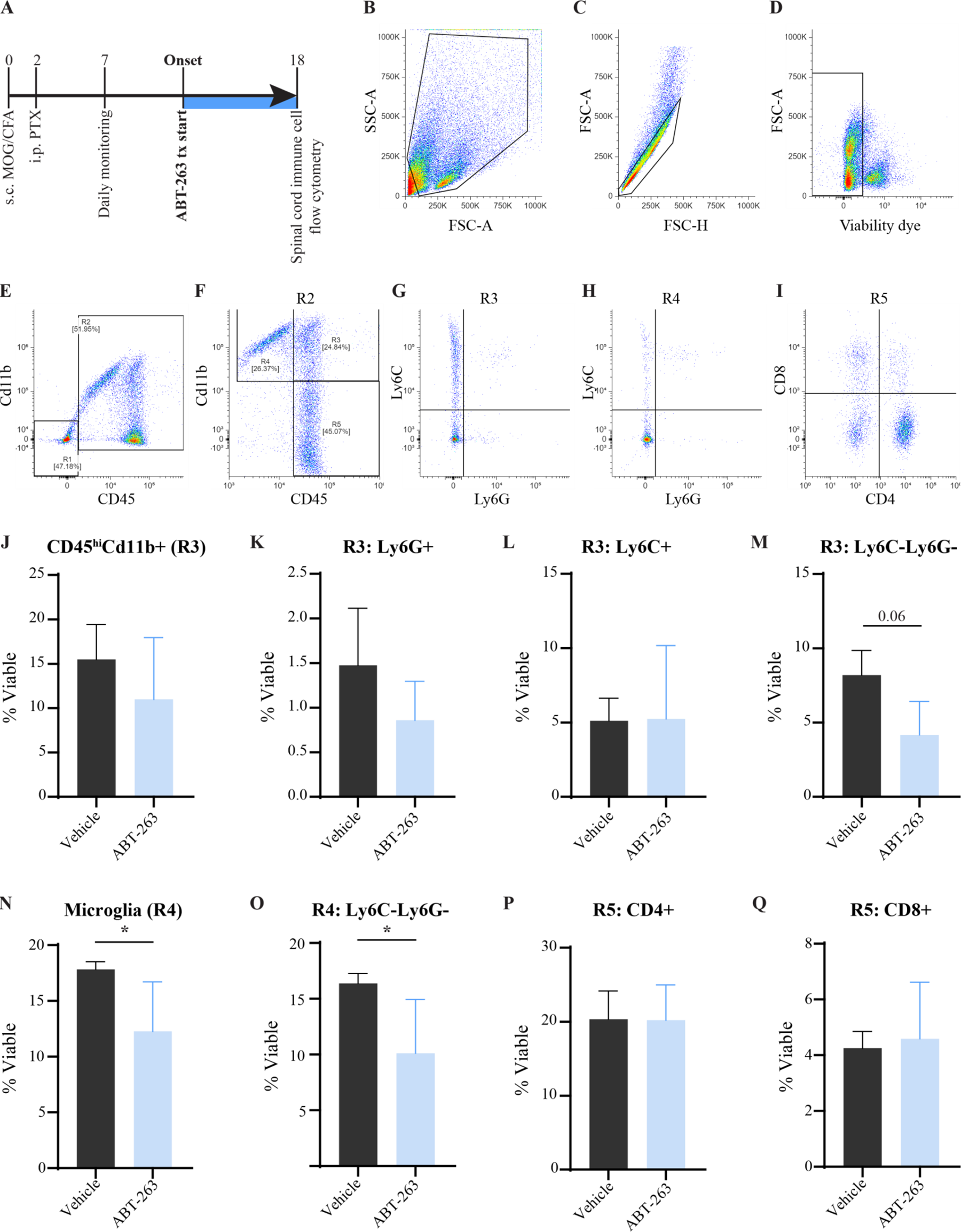
ABT-263 treatment decreases proportion of microglia in spinal cord. **A**) Experimental design demonstrating induction of EAE followed by ABT-263 treatment at symptom onset and spinal cord flow cytometry at 18 d.p.i., and at least 4 days post-treatment start. **B-D**) Cell gating parameters for evaluation of viable single cell population. E) Gating strategy for isolating CD45-Cd11b-cells (R1) from CD45+ cells (R2). **F**) Division of R2 into three cell subsets, consisting of CD45 high Cd11b positive innate immune cells (R3), CD45 mid Cd11b positive microglia (R4) and CD45+Cd11b-lymphocytes (R5). **G)** Division of R3 by Ly6C and Ly6G fluorescence to differentiate monocytes/macrophages (Ly6C-Ly6G-) from other innate immune cells (Ly6C+ proinflammatory monocytes, and Ly6G+ neutrophils). **H**) Division of microglial population based on Ly6C and Ly6G positivity demonstrating most cells (>85%) in the R4 population are negative for both. **I**) Division of the R5 lymphocyte population by CD4 and CD8 positivity. **J-M)** Quantification of the percent of viable cells within the R3 population (R3: innate immune cells, J) including division based on Ly6G (K) and Ly6C (L) positivity, and the R3 population negative for both (M), which was the only population from R3 that approached a significant decrease. **N-O**) Evaluation of the microglial subset in vehicle treated and ABT-263 treated mice as a percent of total viable cells, demonstrating a significant decrease in treated mice in both the whole population (N) and the subpopulation excluding cells positive for Ly6C or Ly6G (O). **P-Q)** Analysis of the R5 lymphocyte population (no change, graph not shown) based on subdivision of CD4 (P) and CD8 (Q) positivity showing no change in the number of CD4+ or CD8+ lymphocytes present in the spinal cords. * < 0.05, all comparison’s conducted with one-tailed Welch’s t-test, vehicle n=2, ABT-263 n=4.

### Senolytic treatment significantly improves EAE severity, neuronal survival, visual function, and white matter inflammation

To evaluate potential therapeutic effects of ABT-263 in the EAE model, mice were treated daily with ABT-263 beginning at disease onset (Fig. 5A). EAE motor symptoms were significantly more mild in ABT-263 animals beginning at 2 days post treatment and continuing until experiment end compared to vehicle treated animals, with a trend as well towards improved weight maintenance (Fig. 5B,C). Likewise, visual acuity, a measure of visual system function, was better maintained in treated animals compared to vehicle controls (Fig. 5D). Next, optic nerve sections were assessed by immunohistochemistry to evaluate the extent of inflammation and demyelination in a uniform white matter tract (Fig.5E-K). Here, immunohistochemistry determined that ABT-263 animals exhibited less Hoechst-stained area in the optic nerve suggesting fewer numbers of immune cells present (Fig. 5E-G). This correlated well with decreased IBA1 positive stained area (Fig. 3H). GFAP positive area was also assessed, for which there was no difference between control and ABT-263 treatment (data not shown). Next, fluoromyelin labeling of lipid membranes demonstrated a trend to higher myelin area (Fig. 5I-K). Particularly, it appeared as around 50% of control treated optic nerves demonstrated significantly decreased myelin area, while none of the ABT-263 optic nerves demonstrated significant myelin loss (Fig. 5K). Lastly, neuronal survival was assessed through BRN3A counts in retinal flatmounts to quantify surviving RGCs in vehicle control and ABT-263 EAE mice (Fig. 5L,M). Remarkably, there was a significant RGC survival effect in the ABT-263 condition (Fig. 5N). Together, these data provide evidence that senolytic treatment inhibiting BCL2 family proteins in EAE mice decreases disease severity, improves outcomes related to visual acuity and neuronal survival, and reduces white matter demyelination and inflammation.

**Figure 5.**
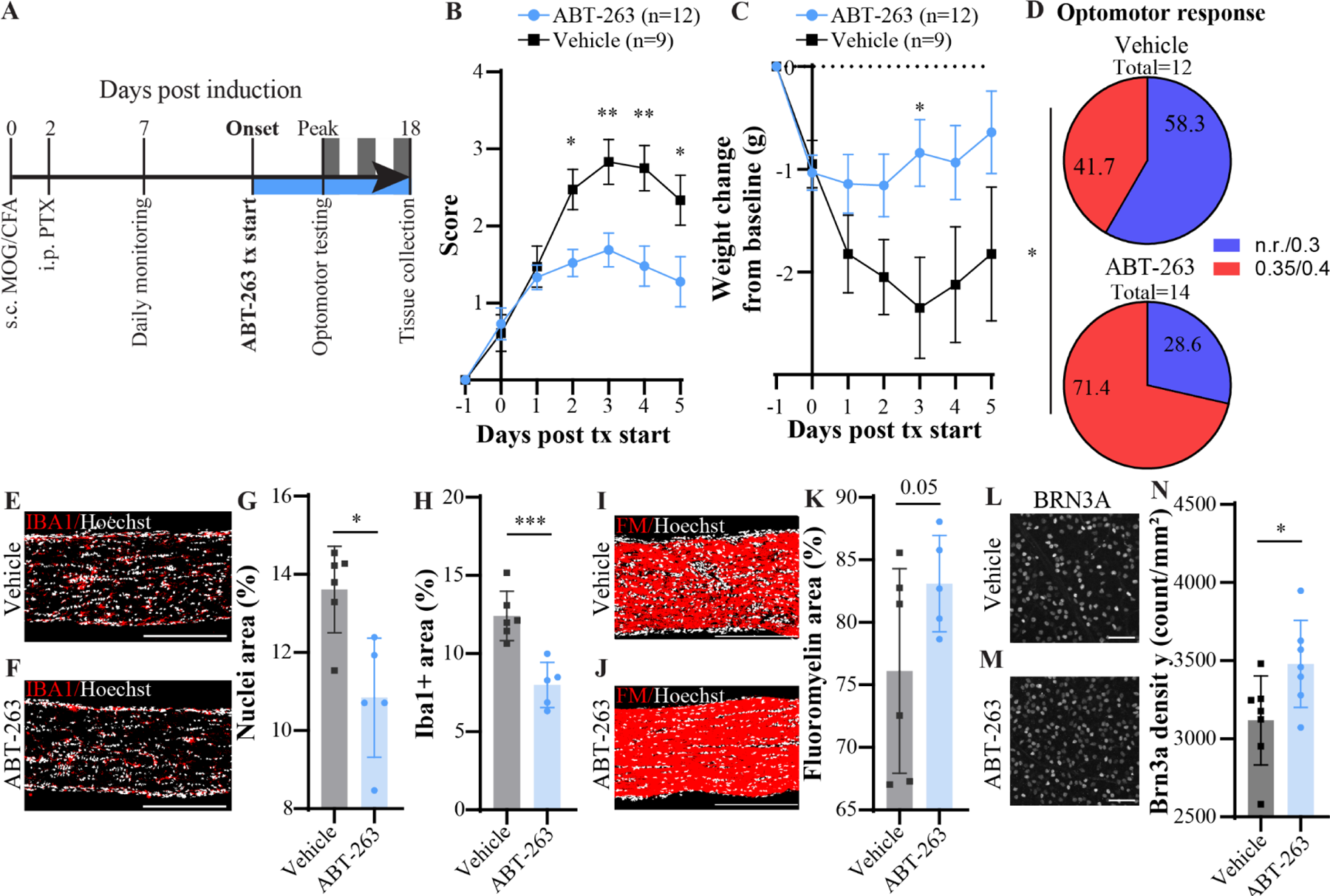
ABT-263 treatment from disease onset reduces severity, inflammation, promotes neuronal survival. **A)** Experimental design demonstrating induction of EAE followed by ABT-263 treatment at symptom onset, optomotor testing at peak disease, and tissue collection at 18 days post induction (d.p.i.). **B)** EAE score from day starting ABT-263 or vehicle treatment, data are combined numbers from two separate EAE cohorts. Bars = SEM. **C)** Weight calculated as difference from baseline of day before symptom onset/treatment start (day 0). Bars = SEM. **D)** Optomotor response data for right and left eyes of EAE mice at peak treated with either vehicle or ABT-263 for at least 3 days prior to evaluation. 3 stripe sizes were tested (0.3 c/d, 0.35 c/d, and 0.4 c/d) and data were binned into two groups no response (n.r.)/0.3 and 0.35/0.4. Number in pie slices represents % of responses in that category. **E,F)** Thresholded images of EAE optic nerves from vehicle treated (E) or ABT-263 treated (F) mice labeled with IBA1 (red) and Hoechst (white). Scale bar = 250 um. **G)** Quantification of the percent area of optic nerve positive for Hoescht dye from thresholded images. Error bars=SD, two-tailed unpaired t-test. **H)** Quantification of the percent area of optic nerve positive for IBA1 immunostaining from thresholded images. Error bars=SD. Two-tailed unpaired t-test. **I,J) T**hresholded images of EAE optic nerves from vehicle treated (I) or ABT-263 treated (J) mice labeled with fluoromyelin (FM-red) and Hoechst (white). Scale bar = 250 um. **K)** Quantification of the percent area of optic nerve positive for Hoescht dye from thresholded images. Error bars=SD, one-tailed Welch’s t-test. **L,M)** Representative images of flatmount retinas labeled with RGC marker BRN3A in vehicle treated (L) and ABT-263 treated (M) EAE mice. Scale = 50um. **N)** Quantification of BRN3A positive cells per area in retinas from vehicle or ABT-263 treated mice. Bars=SD. Two-tailed unpaired t-test.

## Discussion

While there is a consensus that microglia occupy a continuum of states and functions dependent on their intrinsic gene and protein expression, and cues from their local microenvironment, equally there is significant evidence for certain genes consistently dysregulated in microglia across numerous different diseases and in aging^6,10^. It is important to note that many studies on human microglia are often limited by the amount and quality of microglial nuclei, and systematic analysis of microglial gene expression changes compared to mouse gene expression changes in aging and disease finds that microglial signatures comparing across mice and human are not highly similar^2,36^. Still, when considering within humans, microglia demonstrate consistent gene expression changes between aging and neurodegenerative diseases, particularly demonstrating a shared expression profile between microglia from AD patients and MS patients, and between genes downregulated in aging and MS^36^. Equally, within mouse models of aging and neurodegeneration, *Spp1, Ccl2, Ccl3,* and *Ccl4* are frequently upregulated and associated with proinflammatory microglial states^2,37–40^. It seems likely that this gene signature may also be associated with a senescent state among subsets of microglia, as these genes are highly enriched in senescent cells, and senescent microglia have been found in mouse models of Alzheimer’s disease and traumatic brain injury where these genes are upregulated^14,15,22,24,41^. Of note, microglia are essential for myelin maintenance, and aging related changes to autophagic pathways in microglia can interfere with their capacity to properly clear myelin debris^13,38^. Interestingly, ‘regulation of autophagy’ and ‘macroautophagy’ were two of the top enriched GO terms specific to *Bcl2l1+* EAE microglia. In sum, while there are certainly diverse states and gene expression profiles represented in microglia with different specific functions and contributions to disease, the current body of research on microglia in aging and disease create a compelling argument for the presence and contribution of senescent microglia to disease activity in mice and humans^6,40,42^.

Currently, microglia are a major target of interest for development of new MS therapies due to their capacity to perpetuate chronic inflammation and tissue injury in the CNS in the absence of blood-brain-barrier dysfunction and peripheral immune cell invasion^3,5,9,26,43^. For example, the in clinical trial Bruton’s tyrosine kinase (BTK) inhibitors are an exciting new development for MS treatment due to their capacity to pass through the blood-brain-barrier and potentially modulate microglial states therewithin^44^. Indeed, BTK inhibitors may decrease expression of proinflammatory cytokines including TNF, overexpression of which is associated with the proinflammatory, senescent transcriptomic profile seen here in EAE microglia^44^. Targeting microglia is particularly relevant in the case of progressive MS, for which few therapies are currently available and effective at preventing CNS degeneration occurring independent of inflammatory relapses^45^. In MS, reactive microglia particularly contribute to the pathology of chronic active lesions, which are associated with heightened disease progression in patients^3,26,46^. Remarkably, *BCL2L1* positive microglia are almost exclusively present in chronic active lesions, suggesting some relevance to lesion pathology.

The data presented here demonstrate that in EAE, microglia develop a senescent phenotype based on transcriptomics and IHC. However, these senescent microglia constitute only a small proportion of the total microglia, with around 15% of EAE microglia expressing transcripts for *Bcl2l1,* and senolytic treatment resulting in an approximately 5% reduction of spinal cord microglia by flow cytometry and similar reduction in IBA1 positive area in the optic nerve white matter. That said, single-cell RNA-sequencing suggests that the subpopulation of EAE microglia expressing *Bcl2l1* express higher levels of secreted proinflammatory factors. Thus, it is possible that senescent microglia have an outsized effect on inflammation and injury by exacerbating proinflammatory signaling in their local microenvironment and reducing this cell population by ABT-263 treatment broadly reduces CNS glia reactivity and inflammation (Fig. 6). This would be in line with the known function of senescence-associated secretory phenotype, where senescent cells secrete a collection of proinflammatory factors that lead to persistent low-grade tissue inflammation and injury^19^. However, the data presented in this manuscript do not rule out the possibility that other non-microglial senescent cells are being targeted by ABT-263 and contributing to the observed phenotype, as ABT-263 has been demonstrated in other studies to have profound effects on various senescent cell populations (Table 1^22,47–57^).

**Figure 6.**
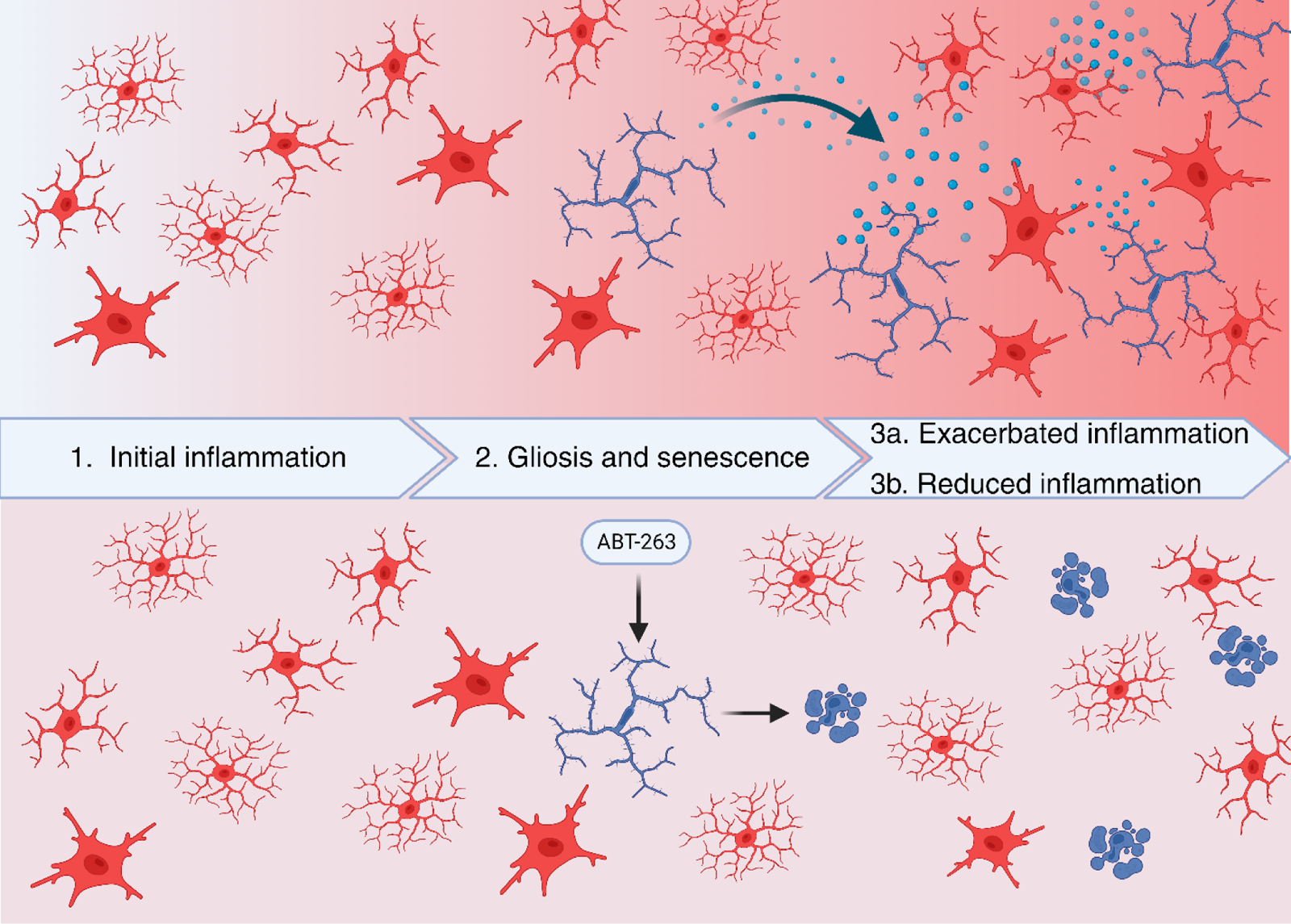
Hypothesis for effectiveness of ABT-263 in EAE. Initial inflammatory insult and injury induces microglia to undergo reactivity and gliosis, and may induce senescence in a subset of microglia (2). Senescent microglia may secrete more proinflammatory factors like Tnf, exacerbating local inflammatory activity (3a). Treatment with ABT-263 may induce apoptosis in the subset of senescent, proinflammatory microglia, reducing the secretion of inflammatory factors into the local milieu, and therefore limiting further inflammatory activity (3b).

**Table 1.** Summary of studies assessing ABT-263 in aging, CNS injury and neurodegenerative diseases^22,36–46^.

| Article | Disease/Model Studied | Implicated cell types | Outcomes of ABT-263 treatment |
| --- | --- | --- | --- |
| Bussian et al, 2018 | Tau dependent neurodegenerative disease | Astrocytes and microglia | Modulated tau aggregation in brains of MAPT<sup>P301S</sup> PS19 mice |
| Acklin et al, 2020 | Chemotherapy induced peripheral neuropathy | Dorsal root ganglion neuronal like cells | Reduced behavioural signs of peripheral neuropathy, p16, and p21 expression. |
| Mehdipour et al, 2021 | Age-related cognitive decline | Microglia | Diminished senescence associated beta-galactosidase in aged mouse brain, reduced area of CD68+ activated microglia stain in hippocampus |
| Paramos-de-Carvalho et al, 2021 | Spinal cord injury | N/A | Improved motor functions following spinal cord injury. Improved myelin sparing, reduced fibrotic scar and inflammation. Decreased secretion of pro-fibrotic and pro-inflammatory factors. |
| Tarantini et al, 2021 | Age-related cognitive decline | N/A | Improved neurovascular coupling, learning and memory. |
| Fatt et al, 2022 | Age-related cognitive decline | Hippocampal neural precursor cells | Depletion of senescent NPCs induced increase in NPC proliferation and neurogenesis. Increased adult-born hippocampal neuron numbers and improved spatial memory. |
| Torres et al, 2022 | Motor neuron disease/degeneration | N/A | hSOD1-G93A transgenic mice show increase of glia with high p16 and p21, but no increase in senescence associated beta galactosidase activity, and ABT-263 treatment did not alter disease progression. |
| Budamagunta et al, 2023 | Age-related cognitive decline | N/A | Rescued memory, preserved blood-brain-barrier integrity, decreased plasma inflammatory cytokine profile, decreased expression of immune markers and increased expression of synaptic markers in the dentate gyrus. Negative regulation of genes related to apoptosis and microglial activation. |
| Lu et al, 2023 | Acute brain ischemia | Astrocytes and cerebral endothelial cells | Improved neurological severity score following brain ischaemic injury, improved locomotor activity, less weight loss. Reduced senescence of astrocytes and endothelial cells. |
| Ahire et al, 2023 | Chemotherapy induced cognitive decline | Endothelial cells | Depletion of senescent endothelial cells, improved neurovascular coupling responses, blood-brain-barrier integrity, and cognitive performance. |
| Gulej et al, 2023 | Radiation therapy induced cognitive decline | Endothelial cells | Depletion of senescent endothelial cells, decreased blood-brain-barrier permeability. |
| Faakye et al, 2023 | Age-related cerebral microhemorrhages | Brain vasculature | Delayed onset, decreased incidence, and severity of cerebral microhemorrhages. |

While these data suggest ABT-263 may be a viable approach to modulate microglia states in MS, further investigation is needed before reaching any clinical application. Assessment of ABT-263 in other EAE models, particularly those that evoke a more progressive disease course would be integral to understanding whether depletion of senescent glia truly ameliorates progressive tissue injury. Additionally, using such a model would enable more efficient evaluation for later treatment start and evaluation of the outcomes of such, better reflecting the treatment intervention timeline for progressive MS patients who may have lived with the disease for many years and accumulated substantial disability and disease activity by the time they receive a therapeutic intervention. Indeed, better characterization as well of senescent microglial states in human MS patient tissue would further solidify these findings. Finally, ABT-263 is a single senolytic, but many exist that may prove more effective in the clinic. For instance, the senolytic drug combination dasatinib and quercetin (D+Q) is already in clinical trials for Alzheimer’s disease, and it is feasible that if successful in AD, this combination therapy may also have off-label success in MS given what seems to be a common feature to many diseases, that senescent glia perpetuate disease activity^58^.

## Acknowledgements

S.S.D. received funding from the Canadian Institutes for Health Research (CIHR) Vanier Canada Graduate Scholarship. E.M.-L.H. is funded by the Fonds de Recherche du Quebec-Santé Master’s Training Scholarship. A.Z. is funded by the Fonds de Recherche du Quebec-Santé PhD Training Scholarship and MS Canada. A.E.F. receives funding from the CIHR and MS Canada. We would like to acknowledge the essential technical support from Thomas Stroh and the Montreal Neurological Institute (MNI) Microscopy Core Facility, and Julien Sirois and the MNI Flow Cytometry Core Facility. We would also like to thank Steve Lacroix for providing advice and a protocol for immune cell flow cytometry.

## Materials and methods

### Animal use approval

All *in vivo* animal experiments were performed using 6-8 week old C57Bl6 mice and experiments received ethical approval according to the guidelines set by the Canadian Council on Animal Care (CCAC).

### Single-cell RNA sequencing analysis

EAE RNA-sequencing data was obtained from the processed files published in Fournier et al. 2023, identified as microglia according to original publication annotations, pseudo bulked and analysed for differential expression following the standard protocol advised in Seurat v.5 documentation^23,59^. Upregulated genes according to differential expression analysis were input for KEGG pathway enrichment using the cluster Profiler package in R^60^. Microglia were subset based on EAE annotation, and each EAE microglial cell was annotated to reflect Bcl2l1 transcript presence or absence (counts > 0, counts = 0). Single-cell differential gene expression was run on EAE microglia grouped by Bcl2l1 positivity and visualized in R. Pathway analysis was run using the goEnrich function from clusterProfiler. Human single-nucleus datasets were obtained from Schirmer et al 2019 and Absinta et al 2022 with original annotations^25,26^. Cluster 7 from the immune subset in the Absinta et al 2022 dataset was used in downstream analyses due to high presence of microglial transcripts and was re-clustered to improve UMAP visualization. Percent positive cells for *BCL2L1* was determined in R for each lesion pathology in both datasets. Due to higher number of cells in Absinta dataset, this dataset was used for downstream differential expression between *BCL2L1* positive and negative cells, and genes were individually visualized with the VlnPlot function in Seurat.

### Experimental autoimmune encephalomyelitis

Active induction of MOG-EAE was achieved as previously described^61–63^. Briefly, 8-week-old female C57Bl6 mice received subcutaneous MOG peptide emulsified in complete Freund’s adjuvant (CFA) in 100 uL volume (50 uL injected bilaterally per mouse). Two days later, mice received intraperitoneal injection of 400 ng pertussis toxin. Mice were monitored daily in the morning for weight change and motor symptom development from 7 days post induction until experiment end. At symptom onset, ABT-263 or vehicle control was administered by intraperitoneal injection daily at 1.5 mg/kg, volume matched for controls based on weight. ABT-263 was dissolved in vehicle solution containing 45% PEG-400, 45% DDH20, and 10% DMSO to a final concentration of 0.5 ug/mL.

### Spinal cord immune cell isolation

Mice were anesthetized, then intracardially perfused with ice cold Hanks Balanced Salt Solution (HBSS w/o Ca^2+^ and Mg^2+^, herein HBSS), followed by decapacitation. Spinal cords were carefully isolated from the spine and immediately transferred into 1X HBSS with 20 mM HEPES. Spinal cord tissue was carefully transferred to a glass homogenizer containing 1 mL of prewarmed enzyme mix (1X HBSS, DNase I 0.025 U/mL, TLCK 0.1ug/mL, HEPES 20 mM, Collagenase type IV 2.5 mg/mL and Elastase 1U/mL) and gently homogenized fifteen times up and down. The tissue and enzyme mix was transferred into a tube and the homogenizer was rinsed twice with enzyme mix that was added to the same tube. Tissue and enzyme mix tubes were incubated for 30 minutes at 37°C, with a homogenization step conducted after 15 minutes of incubation using a P1000 pipette to pipette the tissue and enzyme mix up and down ten times before incubating for the last 15 minutes. After, tubes were spun down at 300g for 3 minutes, resuspended in the pre-existing supernatant, and filtered through a 70 um cell strainer into a new tube. 10 mL of room-temperature wash buffer (1X HBSS,20 mM HEPES, 2 mM EDTA) was added to this mix and the tubes were immediately centrifuged at 300g for 15 minutes. Percoll gradient solutions were prepared in HBSS for 37% and 70%. After centrifugation, supernatant was discarded and the pellet was resuspended in 5 mL of 37% Percoll. 5 mL of the 70% Percoll was carefully underlaid the 37% Percoll/cell suspension, and tubes were centrifuged for 20 minutes at 1000g. Myelin debris was carefully removed from the top layer of the Percoll gradient, and ∼3 mL of the 30-70% interphase was collected into a clean tube and diluted 3X with wash buffer. Tubes were centrifuged at 450g for 10 minutes at 4°C, then the cell pellet was resuspended in 300 uL of cold FACS wash buffer (1X HBSS, 0.5% fetal bovine serum (FBS)) and counted.

### Flow cytometry

Cells were washed twice with cold FACS wash buffer, then blocked with Fc block antibody (4 uL / 100k cells) on ice for 15 minutes. Surface staining mix was prepared with fluorophore conjugated antibodies against Cd11b, CD45, Ly6C, Ly6G, CD4, CD8, and Zombie Aqua for cell viability. Compensation controls were prepared on beads, while FMO controls were prepared on separate cell aliquots. Cells were incubated in a plate with 50 uL of surface staining mix at 4°C in the dark for 30 minutes, then washed three times with cold FACS wash buffer. For flow cytometry settings and parameters, voltages were setup according to optimal PMT sensitivity using the peak 2 (Spherotech) voltration technique described previously by Maeker and Trotter^64^. Compensation control was performed using Ultracomp beads (Thermo Fisher) using optimal antibody concentration determined by titration. All data was acquired on Attune NxT (Thermo-Fisher). Analysis was conducted on all samples in floreada.io in order to obtain percent of viable population of relevant immune cell populations, following similar published gating methods^65,66^.

### Optomotor response

Optomotor response was tested as previously described, in a custom-built drum with striped paper sheets rotated around the animal at 2 rpm in standard lighting^62,67^. Stripe sizes tested included 0.3, 0.35, and 0.4 cycles/degree (c/d). Animals were acclimated to the apparatus for 10 minutes per animal at least one day prior to collecting testing data. Animals that were unable to balance themselves on the platform were excluded from testing. Video data was analysed blinded to mouse identity and condition. Clockwise and counterclockwise responses were measured and compared as a group, such that each animal contributes two datapoints (one result per direction), since each eye is differentially affected in the disease course and contributes the clockwise vs counterclockwise response. Data were aggregated as 0.4 and 0.35, and 0.3 or n.r., as 100% of responses of healthy mice are recorded at 0.35 or 0.4 c/d, thus a response below this threshold indicates decreased visual acuity.

### Tissue collection and processing

Mice were anesthetized then intracardially perfused sequentially with ice-cold PBS and 4% paraformaldehyde (PFA). Eyes, optic nerves, and spinal cords were carefully dissected out and post-fixed in 4% PFA overnight, before washing with PBS. For tissue sectioning, tissues were cryoprotected in 30% sucrose followed by mounting and flash freezing in optimal cutting reagent (OCT). A cryostat was used to collect 12 um sections on slides.

### Immunohistochemistry

For retinal flatmounts, retinae were washed 3x in 1X phosphate buffer solution (PBS) then blocked overnight in 5% donkey serum and 0.2% Triton-X (blocking solution). After blocking, retinas were incubated in primary mouse anti-Brn3a antibody at 1:500 in 1% donkey serum and 0.2% Triton-X (staining solution) for 3 days at 4°C. Afterwards, tissue was washed 3X with PBS then incubated in secondary antibodies, AF568 donkey anti-mouse at 1:500 and Hoechst at 1:1000, in staining solution over night at 4°C. Tissue was then washed 3X with PBS then mounted on slides with coverslips and Fluoromount mounting reagent. For tissue sections, sections were rehydrated with PBS and washed 3X with PBS. Then, sections were blocked with blocking solution for 1 hour at room temperature, followed by primary antibodies overnight in staining solution at 4°C. Next, tissue was washed 3X with PBS and incubated with secondary antibodies in staining solution at room temperature for 1 to 3 hours. Finally, sections were washed 3X with PBS and coverslips were mounted on slides with Fluoromount mounting reagent. Senescence associated beta galactosidase fluorescence was conducted using the SpiderGAL kit according to manufacturer’s instructions prior to IHC.

### Imaging and analysis

Images were acquired on an LSM 880 confocal microscope using z-stacking and tiling functions to image the entire length of optic nerve sections with the same laser intensity and settings across all samples. Images were imported into ImageJ and regions of interest were drawn manually to isolate the optic nerve. Then, default threshold parameters were used in ImageJ to generate a thresholded image, and %area covered within the optic nerve region was measured in ImageJ.

